# *In-vitro* and *in-vivo* Characterization of a Multi-Stage Enzyme-Responsive Nanoparticle-in-Microgel Pulmonary Drug Delivery System

**DOI:** 10.1101/650911

**Authors:** Joscelyn C. Mejías, Krishnendu Roy

## Abstract

Although the lung is an obvious target for site-specific delivery of many therapeutics for respiratory airway diseases such as asthma, COPD, and cystic fibrosis, novel strategies are needed to avoid key physiologic barriers for efficient delivery and controlled release of therapeutics to the lungs. Specifically, deposition into the deep lung requires particles with a 1-5 µm aerodynamic diameter; however, particles with a geometric diameter less than 6 µm are rapidly cleared by alveolar macrophages. Additionally, epithelial, endothelial, and fibroblast cells prefer smaller (< 300 nm) nanoparticles for efficient endocytosis. Here we address these contradictory design requirements by using a nanoparticle-inside-microgel system (Nano-in-Microgel). Using an improved maleimide-thiol based Michael Addition during (water-in-oil) Emulsion (MADE) method, we fabricated both trypsin-responsive and neutrophil elastase-responsive polymeric Nano-in-Microgel to show the versatility of the system in easily exchanging enzyme-responsive crosslinkers for disease-specific proteases. By varying the initial macromer concentration, from 20-50 % w/v, the size distribution means ranged from 4-8 µm, enzymatic degradation of the microgels is within 30 minutes, and *in vitro* macrophage phagocytosis is lower for the higher % w/v. We further demonstrated that *in vivo* lung delivery of the multi-stage carriers through the pulmonary route yields particle retention up to several hours and followed by clearance within in naïve mice. Our results provide a further understanding of how enzymatically-degradable multi-stage polymeric carriers can be used for pulmonary drug delivery.

**Graphical Abstract:** 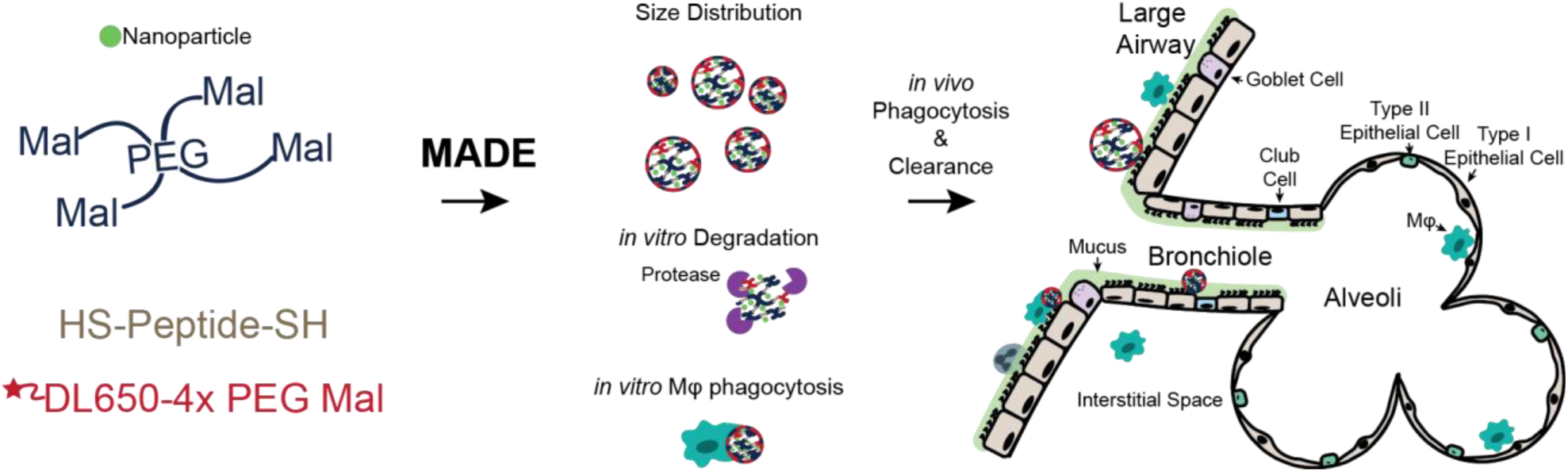

## Introduction

Asthma, chronic obstructive pulmonary disorder (COPD), and cystic fibrosis (CF) are chronic inflammatory diseases characterized by an influx of innate immune cells to the lungs, including macrophages[1–3], neutrophils[4, 5], and eosinophils[2]. These cells impose a heavy enzymatic burden through proteases such as matrix metalloproteinases and elastases that promote significant tissue damage. Pulmonary delivery is a potentially non-invasive administration route for treating these chronic inflammatory disorders; however, the anatomy and physiology of the lung present significant design challenges for achieving efficient and effective drug delivery. Nanoparticles can inherently provide protection for encapsulated biologic therapeutics (proteins, RNAs) from high levels of exogenous proteases present in lung diseases, but they must be smaller than 300 nm to provide efficient cellular endocytosis[6–8] and transport across the mucosal barrier to target epithelial cells[9–12]. However, nanoparticles alone cannot efficiently aerosolize and deposit within the lung via inhalation; particles with aerodynamic diameters less than 1 µm are predominantly exhaled, while those between 1-5 µm are deposited more efficiently into the deep lung, and those greater than 5 µm are impacted in the mouth or throat[13]. At the same time, alveolar macrophages rapidly phagocytose particles with geometric diameters within the 1-5 µm range[14, 15], making efficient delivery difficult.

Current strategies for addressing this conflicting design issues include the use of porous polymers and swellable microparticles. By lowering the density of the microparticles, large porous particles provide an aerodynamic diameter between 1-5 µm and a geometric diameter greater than 6 µm[16–18] thus increasing residence time after deposition in the lungs. Swellable microparticles in their relaxed, dry state provide the correct aerodynamic size for lung delivery, and swell in the humid lung environments to a size capable of avoiding rapid macrophage clearance[19–22]. However, once these microscale carriers reach the deep lung, transport across mucosal surfaces and intracellular delivery to target cells, remain significant challenges.

The goal of this research was to synthesize and characterize a multi-stage, Nano-in-Microgel pulmonary delivery system for treating chronic inflammatory lung diseases that addresses the following complex design challenges - (1) appropriate aerodynamic size to achieve deep lung delivery, (2) protease-triggered release of encapsulated nanoparticles, and (3) avoiding rapid alveolar macrophage clearance. Nano-in-Micro particles have been previously reported for intestinal[23, 24] and lung delivery[19–22]. Our group have previously reported an *in vitro* proof-of-concept Nano-in-Microgel formulation using a trypsin-responsive crosslinker with 60% w/v 4-arm-PEG acrylate[19]. Also, poly(ethylene glycol) based hydrogels with protease responsive backbones have been used to respond and degrade to the cellular microenvironment[25–27].

Here we report detailed *in vivo* characterization of a neutrophil elastase responsive Nano-in-Microgel formulation and evaluate whether such a delivery system can achieve the three critical design criteria outlined above. The neutrophil elastase responsive peptide was chosen as this protease is well known to be involved in various acute and chronic pulmonary inflammatory diseases[4, 28–30]. Specifically we hypothesized that microgels crosslinked with elastase-responsive peptides would rapidly degrade under clinically-relevant conditions – i.e. in elastase-rich bronchial fluid from cystic fibrosis patients (design criterion 2), and when delivered *in vivo* in naïve animals they will reach the deep lung space (i.e. show appropriate aerodynamic properties, design criterion 1) without being rapidly cleared by alveolar macrophages (design criterion 3).

Rapid alveolar clearance has been shown to occur within 30 minutes of microparticle administration to the lung in animal models and hinders drug delivery[16]. Here we evaluated the impact of different starting polymer concentrations on avoiding rapid macrophage clearance using two different peptide crosslinkers, trypsin-responsive CGRGGC, and neutrophil elastase-responsive CGAAPVRGGGGC (adapted from [31]). We also developed an improved Michael-Addition-During-(water-in-oil)-Emulsion (MADE) method which provides faster reaction kinetics during fabrication while avoiding degradative conditions such as UV light, high temperatures, and organic solvents that could denature biologic drugs or sensitive therapeutics. We report *in vivo* performance of these Nano-in-Microgels following pulmonary delivery in naïve mice, including addressing their pulmonary biodistribution, cellular uptake, and clearance kinetics. Our results show that multi-stage Nano-in-Microgels are a promising delivery system for efficient intracellular delivery to the lung.

## Materials and Methods

### Microparticle Formulation

Microgels were fabricated using Michael Addition During Water-in-Oil Emulsion (MADE). Briefly, equimolar amounts of a di-sulfhydryl peptide (CGRGGC, CGAAPVRGGGGC; CHI Scientific, Maynard, USA) and a 4-arm PEG Maleimide (Mal, 10kDa, Laysan Bio, Arab, USA) were dissolved separately in PBS to the desired combined concentration (e.g., 20 % w/v is 20 mg of molar ratio thiol:Mal in 100µL of PBS, Supplemental Table 1). To physically entrap the nanoparticles within the PEG network during crosslinking, the nanoparticles were resuspended in the PEG macromere solution. The PEG-nanoparticle-peptide mixture was added to 15 mL of mineral oil (Sigma Aldrich, St. Louis, USA) with 1% v/v surfactant (Span 80/Tween 80, HLB = 5, Sigma Aldrich) and homogenized on a PRO Scientific D Series homogenizer (Oxford, USA) for three minutes at 4000 rpm in a 35-40°C water bath. The emulsion was then left rotating for a minimum of two hours at 37°C. Microgels were removed from the surfactant and residual unreacted material by multiple centrifuge washes in water. To make non-enzyme responsive microgels, a 1 kDa HS-PEG-SH (Creative PEGWorks, Chapel Hill, USA) was used in place of the peptide, and the solution pH lowered to increase the crosslinking time. Microgels were filtered through a 40 µm cell filter before use to prevent clogging of the flow cytometers and the Penn-Century Inc. MicroSprayer (Wyndmoor, USA). Nanoparticles encapsulated include: 0.1 ± 0.012 µm Blue FluoSpheres (ThermoFisher Scientific, Waltham, USA, ex/em 350/440) or 60 nm Carboxylated Dragon Green polystyrene beads (Bang’s Laboratories, Fishers, USA, ex/em 480/520). Microgels were labeled for microscopy or flow cytometry with 0.01 mg of DyLight 650-4xPEG-mal, DyLight 650-mal or DyLight 488-mal (ThermoFisher, dissolved at 10 mg/mL in DMSO). Encapsulation efficiency of Blue FluoSpheres was measured by fluorescence of nanoparticles against a standard curve with a BioTek (Winooski, USA) plate reader.

### Microscopy and Analysis

Fluorescent images were collected either on a Zeiss (Oberkochen, Germany) 710 Laser Scanning Confocal Microscope with a 63X NA-1.4 oil Plan Apochromat objective or on a PerkinElmer (Waltham, USA) UltraVIEW VoX spinning disk confocal microscope with a Hamamatsu C9100-23b back-thinned EM-CCD and Nikon 100x NA-1.45 oil objective. Sizing of swollen particles was performed with the Matlab function “imfindcircles”. Other image processing, linear contrasts, particle-size in oil, calculations, stitching, and three-dimensional reconstruction were performed in Volocity (PerkinElmer).

### Rheometry

Hydrogel storage and loss moduli were measured by dynamic oscillatory strain, and frequency sweeps using an MCR 302 Rheometer (Anton Paar, Graz, Austria) with a 9 mm diameter 2° cone and plate set at 25°C. Bulk hydrogels were made in 4.5 mm diameter silicone isolators, crosslinked, then removed from the isolators and allowed to swell in PBS overnight. One gel per formulation was used to determine the linear viscoelastic range of the hydrogel using a strain amplitude sweep with an angular frequency of 10 rad/s; strain from the linear portion was then used for the oscillatory frequency sweeps. Frequency sweeps were run with the constant strain from ω = 0.1-100 rad/s or until at least six points were linear.

### In-vitro Degradation

Microgels were studied in both the absence and presence of enzyme using time-lapse imaging of a loaded cell trap provided by the Lu lab [32]. The use of a cell trap allows for the imaging of Nano-in-Microgel degradation over time with a high n (n > 100) per run. Once traps were loaded DyLight 650 labeled microgels, enzyme solutions were flown into the trap, and the trap was fluorescently imaged every minute for 30 minutes on a BioTek Lionheart using a 4x objective. Images were stitched and analyzed in Gen 5.02 (BioTek). De-identified CF pooled patient sputum was collected by the Emory Biospecimens Repository under protocols approved by the Institutional Review Boards at Emory University and Georgia Institute of Technology.

### Macrophage Uptake Study

RAW 264.7 Macrophage cells (ATCC, Manassas, USA) were cultured in DMEM medium supplemented with 10% Fetal Bovine Serum (FBS) 100 U/mL/100 µg/mL Penicillin-Streptomycin (HyClone, Logan, USA) in a 37°C incubator with 5% CO_2_. RAW cells were detached by scraping and stained with CellTrace Yellow (ThermoFisher) following manufacturer instructions, then incubated with 20-50% w/v Nano-in-Microgel at a 1:1 ratio in a 96 well V-bottom plate. At 0.5, 1, 2, 6, and 12 hours the plates were moved to 4°C, and the cells stained with Zombie Green (BioLegend, San Diego, USA) and fixed with Cytofix (BD Biosciences, San Jose, USA). Fluorescence was evaluated using a BD LSRFortessa with a high-throughput system.

### Pulmonary Administration and whole organ imaging

All studies were performed according to protocols approved by the IACUC at the Georgia Institute of Technology and all methods were performed in accordance with the relevant guidelines and regulations. C57BL/6 and 8-12 weeks old female Balb/c mice (Jackson Laboratories, Bar Harbor, USA) were maintained in pathogen-free facilities, given sterile water, and food (low alfalfa if used for imaging) ad libitum. Using a 1A-1C MicroSprayer Aerosolizer and FMJ-250 high pressure syringe assembly (Penn-Century, Inc) for aerosolized delivery, mice were anesthetized with a Ketamine/Xylazine 80-100 mg/kg/ 10-20 mg/kg cocktail, positioned on an intubation table, endotracheally intubated with a 1.22 mm ET Tube (Hallowell EMC, Pittsfield, USA), and swollen Nano-in-Microgel formulations (1e6 microgels in 50 µL) were injected. If there was injection failure, the mouse was marked for follow up and removed from data sets; all samples removed are marked within the relevant supplemental figures. For oropharyngeal aspirations, mice were anesthetized with isoflurane, placed on an intubation table, then the tongue was pulled and held with atraumatic forceps as the formulation was administered into the mouth, and the nose gently covered until the liquid was inhaled. Within experiments, the concentration of microgels was equal between formulations. After sacrificing the mice with sodium pentobarbital, perfusing the lungs with PBS through the right ventricle then excising the lungs, the fluorescence intensity of the excised lungs was measured by imaging via IVIS Spectrum µCT (Perkin Elmer).

### Flow Cytometry to study *in vivo* cellular targeting and clearance

Single cell suspensions were prepared using a gentleMACS tissue dissociator (Miltenyi Biotec, Bergisch Gladbach, Germany) with 1 mg/mL of Collagenase D (Roche, Basel, Switzerland) and 0.1 mg/mL DNase I (Roche) in Opti-MEM (ThermoFisher), then straining through a 40 µm cell strainer. Red blood cells (RBC) were lysed using 1x RBC Lysis Buffer (eBioscience, San Diego, USA). The single cell suspension was then washed and stained with Zombie Aqua (BioLegend, San Diego, USA), incubated with Fc Block (BioLegend) then with fluorochrome-conjugated antibodies (BioLegend unless noted; clone used), and fixed with Cytofix (BD Bioscience). Antibodies used were: Ly-6G-BV711 (1A8), CD45-BV605 (30-F11), SiglecF-PE (BD Bioscience; E50-2440), CD11c-APC/Fire750 (N418), F4/80-PE/Dazzle™ 594 (BM8), CD11b-PE/Cy7 (M1/70). Cells were collected on an untreated littermate to set the gates with noise below 1% by FMOs (fluorescence minus one). The gating scheme is illustrated in Supplemental Figure 1. Data were acquired on a BD LSRFortessa flow cytometer; compensation and data analyses were performed using FlowJo (TreeStar, Ashland USA).

### Statistical Analysis

Data was analyzed using GraphPad Prism (San Diego, CA) with a Shapiro-Wilk test for normality (α=0.05). A one-way Anova (α=0.01) with post-hoc Tukey’s test with an adjusted P value for multiple comparisons, a two-way Anova (α = 0.01) with Sidak post hoc analysis was used with data that passed Shapiro-Wilk. Where applicable, a Mann-Whitney (α = 0.01) or Kruskal-Wallis (α = 0.01) with Dunn’s test was used for data that failed Shapiro-Wilk.

## Results and Discussion

### Design and Synthesis of Nano-in-Microgel Particles for aerodynamic and geometric size

Microgel formulations were made at 20, 30, 40, 50% w/v starting polymer concentration to assess the effect on microgel size, degradation, and macrophage phagocytosis. 4-arm PEG Maleimide was crosslinked with a trypsin-responsive (CGRGGC) or neutrophil elastase-responsive (CGAAPVRGGGGC) peptide backbone via crosslinking of the thiols on the cysteine residues. A maleimide-thiol reaction was chosen for the Michael Addition During water-in-oil Emulsion because of the faster kinetics relative to an acrylate-thiol[33]. By using both trypsin- and elastase-degradable crosslinker, we show the versatility of the multi-stage system, whereby the crosslinker can be exchanged for different disease-related protease responsive sequences. These peptides allow for the controlled degradation of the microgels by proteases to release their nanoparticle cargo, schematic representation in Figure 1A. To fluorescently label the microgel for imaging and flow cytometry, a DyLight 650-4x-PEG-Maleimide was added to the 4x-PEG-Maleimide precursor solution.

**Figure 1:**
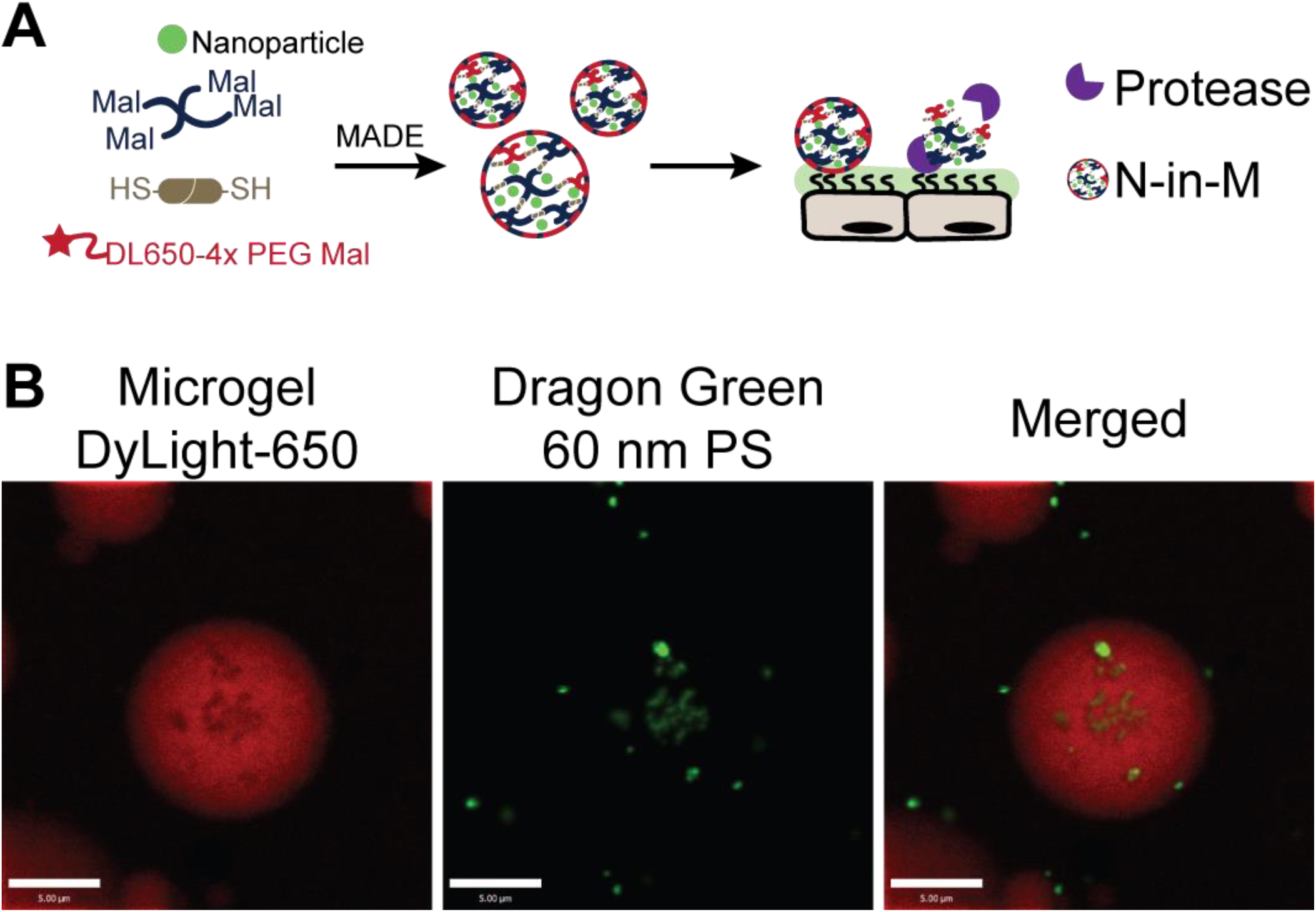
Synthesis of the Nanoparticle-in-Microgel (N-in-M) System. (A) Schematic of the synthesis and degradation process for the Nano-in-Microgel formulation. To fabricate the Nano-in-Microgels, a Michael Addition reaction During water-in-oil Emulsion (MADE) was used. 4-arm PEG Maleimide and DyLight 650 4-arm PEG Mal is crosslinked by a protease-responsive di-thiol peptide in emulsion. During microgel fabrication, nanoparticles are physically entrapped into the microgels. Following delivery, protease dependent degradation at tissue or disease sites leads to nanoparticle release (B) Representative image of a 20% trypsin-responsive DyLight 650 labeled microgel encapsulating 60 nm Dragon Green polystyrene beads. Scale bars = 5 µm. Contrast linearly adjusted for clarity.

Representative images of a DyLight-650 labeled microgel with 60 nm carboxylated polystyrene beads can be seen in Figure 1B. Carboxylated beads (60 nm) can be seen in more than 89% of the microgels by flow cytometry, Supplemental Figure 2. Encapsulation efficiency of the 100 nm Blue FluoSpheres for each formulation was between 30-40% with total nanoparticles encapsulated (∼1e11) in the microgels within the same order of magnitude as theoretical maximum (3.6e11), Supplemental Table 2.

The mean geometric diameter of the microgels for each starting polymer concentration were between 4.5-8.5 µm, Figure 2A. Mean size generally showed an increasing trend with the increase of starting polymer concentrations; however, they were not statistically different. The lowest size is limited by the imaging technique used for the measurement. 60% w/v microgels were not included in the studies as their yield was low and the viscosity of the solution was difficult to transfer to the homogenizer efficiently.

**Figure 2:**
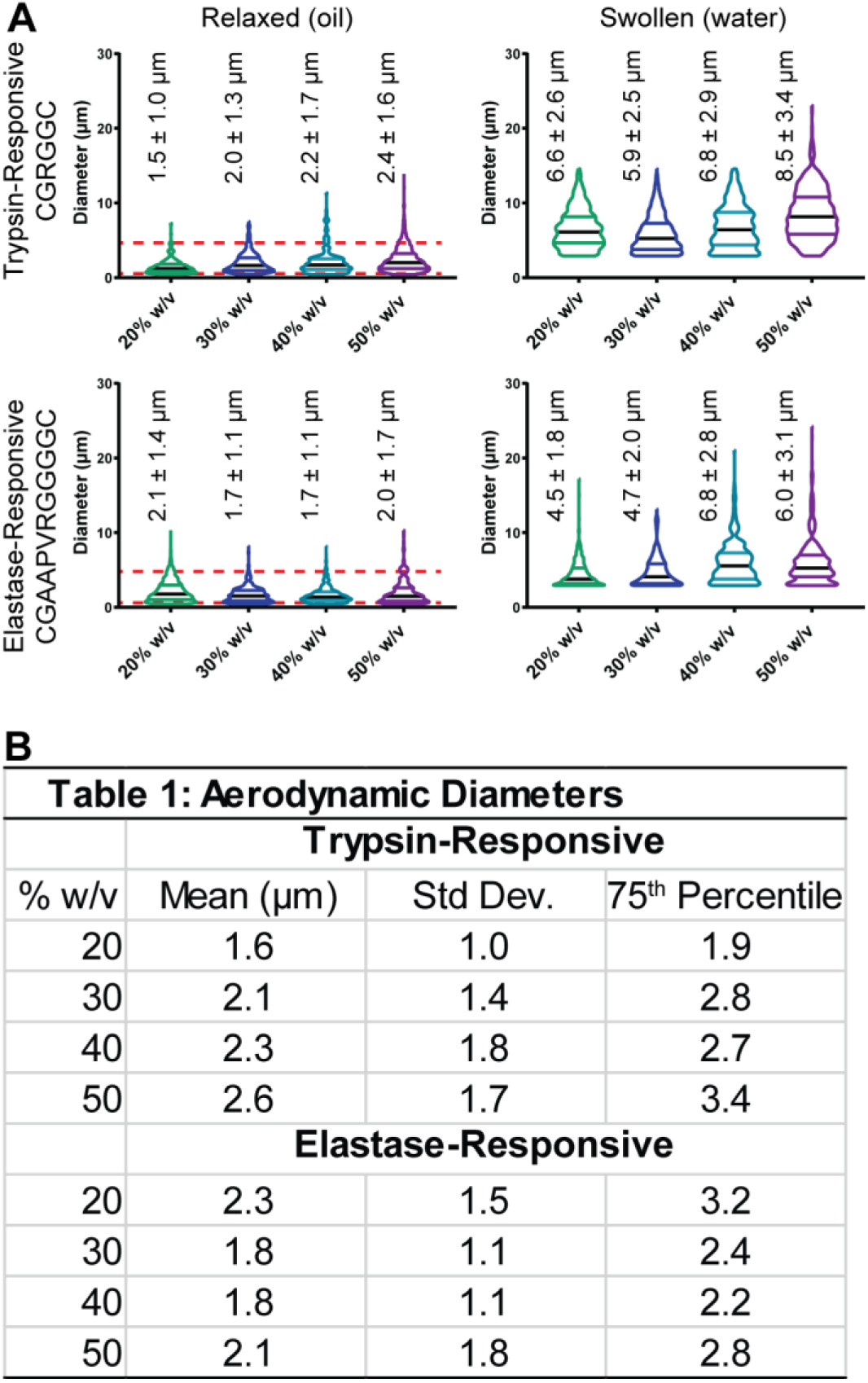
Size distribution of the trypsin- and elastase-responsive Nano-in-Microgel formulations in relaxed vs swollen states. (A) Microgels fabricated with a trypsin responsive (CGRGGC) peptide sequence or a neutrophil-elastase responsive (CGAAPVRGGGGC) peptide with varying starting polymer concentrations 20% (green), 30% (blue), 40% (cyan), 50% (purple) were sized via microscopy in their relaxed polymer state (in oil) and swollen state (in water). The dashed red lines on the relaxed state graphs represent the theoretical range of geometric diameters that correspond to 0.5-5 µm aerodynamic range, which is thought to be appropriate for deep lung delivery [13, 14]. These were calculated by the relationship: D_a_ = D_g_*ρ^0.5^ for Da = 0.5 and 5. Each violin plot is from max to min with median (black line) and quartiles labeled, n > 200 for each measurement. (B) The table shows the mean, standard deviation (Std Dev.) and 75^th^ percentile aerodynamic diameters for the trypsin- and elastase-responsive microgels from 20-50% w/v starting polymer concentrations, calculated from the geometric diameters using D_a_ = D_g_*ρ^0.5^.

Aerodynamic diameter (D_a_) is dependent on the geometric diameter (D_g_), shape factor (1 for spheres) and density of the particle through the following relationship: D_a_ = D_g_*ρ^0.5^ [14]. Using the density of PEG as 1.13 g/mL, aerodynamic diameters of various microgel formulations were calculated and shown in Table 1 (Figure 2B). Aerodynamic diameters of 0.5 to 5 µm are most suitable for deep lung deposition [13, 14]; which corresponds to a theoretical geometric size range of 0.94 and 4.7 µm for PEG based particles. This range is represented by dashed red lines on Figure 2B and suggests that the design criteria for optimal aerodynamic diameter for deep lung delivery is met. In the future, the actual aerodynamic diameter measured by a next generation impactor, after developing a dry powder formulation, would be necessary to account for microgel aggregation, density, and shape factor.

### More than 95% of Microgels degrade within 30 minutes in vitro

In order to confirm the design criteria for proteolytic degradation of the microgels by the incorporated peptide backbone, Nano-in-Microgels were loaded into microfluidic, PDMS single cell traps [32] then exposed to the relevant proteases and imaged over time (Figure 3A). Trypsin-responsive microgels were exposed to 0.01 mg/mL of trypsin (≥ 90 u/mL). For all formulations, 95% of the microgels degraded within 20 minutes. These studies were repeated with the elastase-responsive microgels, which were exposed to pooled sputum from cystic fibrosis patients (courtesy of the Emory Cystic Fibrosis Center, Atlanta, GA). For all elastase-responsive microgel formulations, 95% of the microgels degraded within 25 minutes of exposure to the sputum. It is possible that the protease-responsive crosslinkers in these microgels could non-specifically degraded by other proteases present in the disease state; e.g., matriptase for the trypsin-responsive peptide or cathepsin S for the elastase-responsive peptide (identified using PACMANs[34]), and this *in vivo* degradation rate in disease states could be faster – an aspect we plan to evaluate in future studies. Non-enzyme degradable 20% w/v microgels were fabricated by crosslinking with a dithiol PEG and exposed to 0.01 mg/mL of trypsin (≥ 900 u/mL) as a control to show the peptide crosslinker is necessary for enzyme-responsive degradation. The fluorescence change of the non-enzymatically degradable PEG microgels followed similar trend as several sets of trypsin and elastase crosslinked microgels that were imaged without exposure to the enzyme to confirm limited bleaching of the microgels; the mean at 25 minutes is 91%, Figure 3B – indicating minimal background degradation.

**Figure 3:**
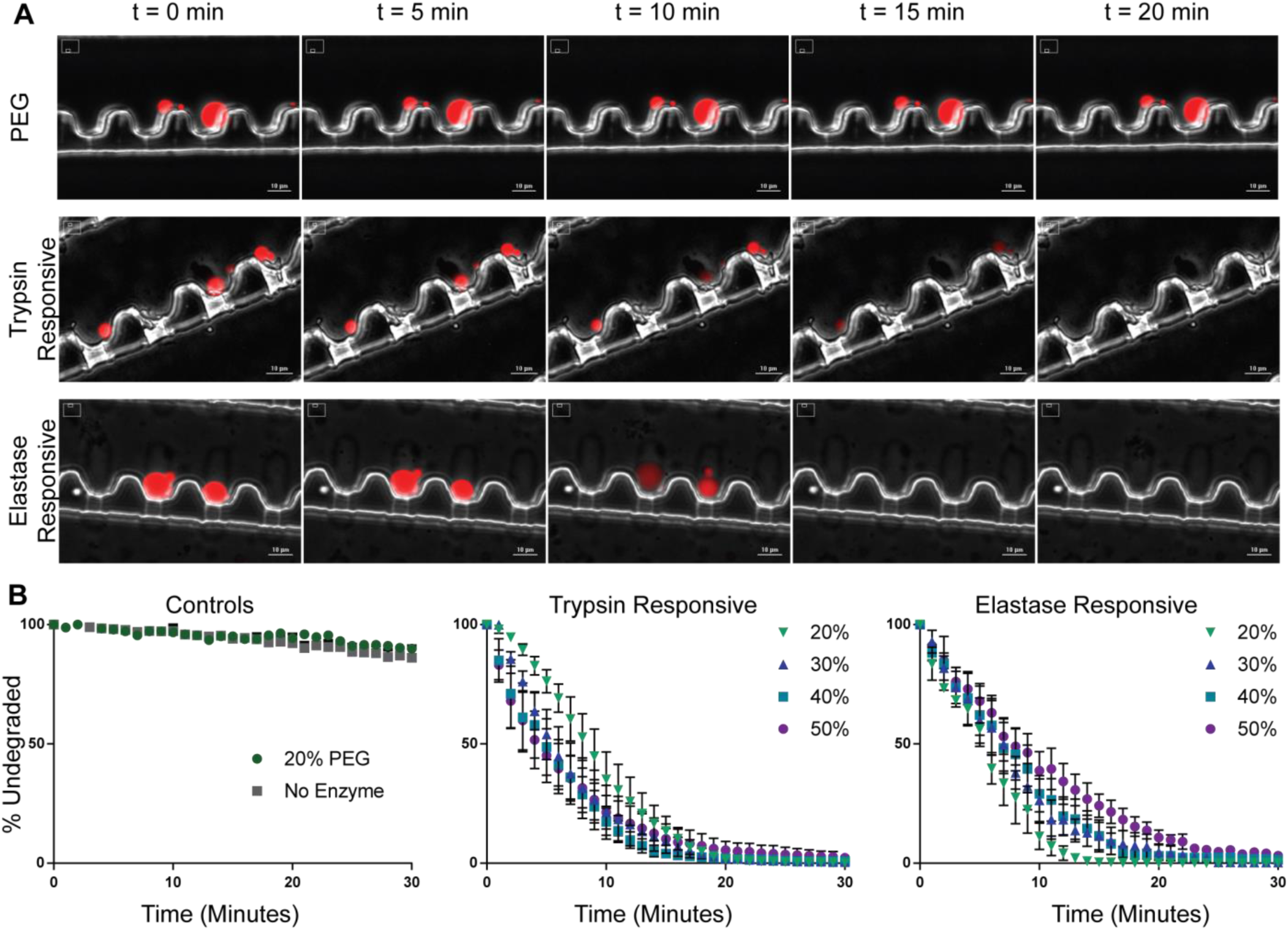
Degradation of trypsin- and elastase-responsive microgels by their respective proteases. (A) Representative phase contrast, Cy5 overlaid images of cell traps at 0, 5, 10, 15, and 20 minutes holding microgels crosslinked with PEG (non-protease responsive), trypsin-, or elastase-responsive microgels. Whole cell traps were imaged before protease addition and once every minute for 30 minutes after addition. PEG was imaged with and without exposure to 0.01 mg/mL of trypsin to measure photobleaching and show the necessity of the protease-responsive peptide for the microgel degradation. Trypsin-responsive microgels were exposed to 0.01 mg/mL trypsin and elastase-responsive were exposed to pooled CF patient-derived airway supernatant; scale bar = 10 µm. (B) Mean ± SEM plots of the percent of non-degraded microgels remaining in the trap over 30 minutes for PEG microgels (N = 5), trypsin- (N = 5) and elastase- (N = 3) responsive microgels. Trypsin- and elastase-responsive microgels were tested for each 20-50% w/v starting polymer concentration with 95% of trypsin-responsive microgels degrading in 20 minutes and 95% of elastase-responsive degrading in 25 minutes.

Interestingly, we did not observe a difference in degradation when varying polymer concentrations. This could be due to either the limits of the measurement method (1-minute intervals) or the polydispersity of the microgels and overlap in size of the different groups. It is also possible that the enzyme concentration is high and the kinetics of degradation is much faster than the increase in number of additional crosslinks. It should be noted that the release of the 100 nm nanoparticles should follow the degradation curve, since the theoretical pore size calculated[35] is smaller than the size of the nanoparticles (10-20 nm) i.e., the nanoparticles can only be released as the microgels degrade, Supplemental Table 3.

### Bulk hydrogels from 20-50% are relatively soft

Since macrophage phagocytosis (the final design criteria) has been shown to be dependent on size and mechanical properties of the particles [36, 37], we next measured storage (G’) and loss (G’’) moduli of the bulk hydrogels (Figure 4). For the trypsin-responsive crosslinker, the 20% w/v microgels had significantly lower loss modulus than the 40 and 50% w/v ones. Furthermore, both the trypsin and elastase-responsive 20% w/v formulation had lower storage modulus than the 40 and 50% w/v bulk gels. However, across all formulations, the hydrogels are relatively soft (10kPa).

**Figure 4:**
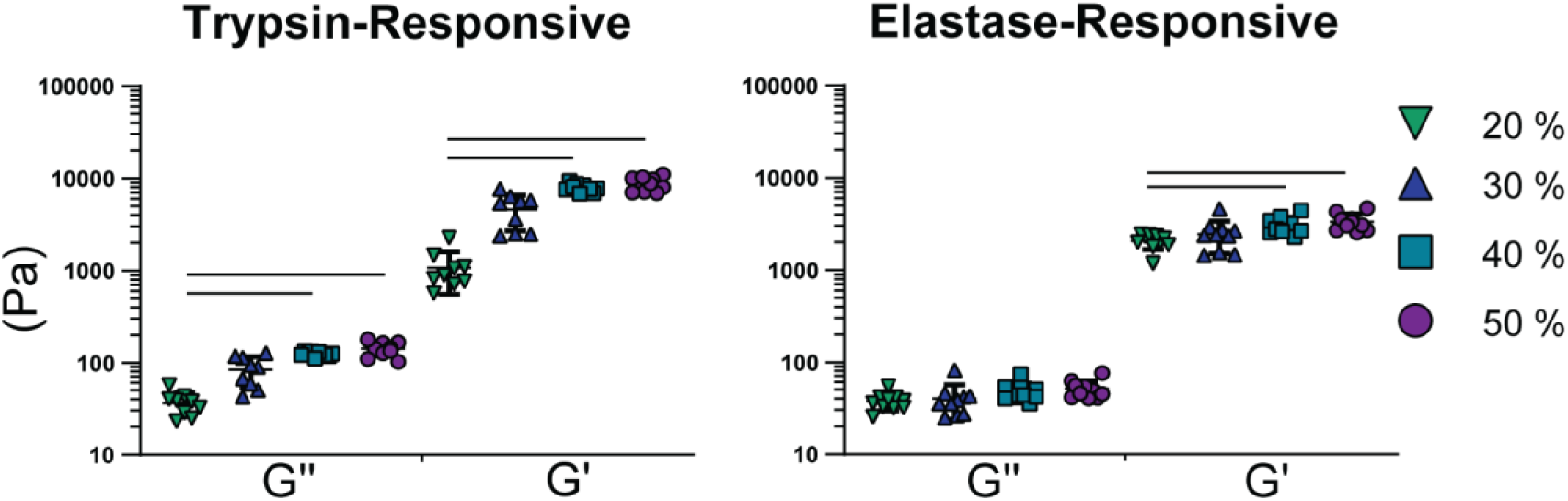
Storage (G’) and loss (G’’) moduli for trypsin- and elastase-responsive crosslinked bulk hydrogels. Trypsin- and elastase-responsive bulk hydrogels with varying starting polymer concentrations from 20-50% w/v (n = 8). Bars within formulation are group mean ± SD. Horizontal bars represent statistical significance, Kruskal-Wallis with Dunn’s multiple comparison test (α < 0.01)

### Higher polymer concentration results in lower phagocytosis of microgels by macrophages in vitro

To study macrophage phagocytosis *in vitro*, DyLight 650-4xPEG mal labeled Nano-in-Microgel formulation (20-50% w/v) loaded with 100 nm Blue FluoSpheres were mixed and incubated with RAW 264.7 cells at a 1:1 cell:particle ratio. At 0.5, 1, 2, 6, and 12 hours of incubation, the cells were fixed and analyzed using flow cytometry for Nano-in-Microgels positive macrophages (double positive DyLight 650 and Blue FluoSpheres) or nanoparticle only macrophages (Blue FluoSpheres), Supplemental Figure 3A. After fixing, a small sample of cells from each treatment group was placed on a slide and imaged on a spinning disc confocal. A representative image (Figure 5A) showed that macrophages appear to phagocytose multiple particles. Over time, the percentage of macrophages that were positive for the Nano-in-Microgels increased for all formulations; however, Nano-in-Microgels with a higher polymer concentrations exhibited lower uptake by macrophages *in vitro*. This trend was observed for both the trypsin- and elastase-responsive microgels, Figure 5B, and suggests that to avoid rapid clearance by phagocytes a higher starting polymer concentration may potentially be helpful *in vivo*. The size and rheological properties of the microgels may have a combined effect resulting in differential phagocytosis. The RAW 264.7 macrophages were not activated with any immune-stimulatory signals, nor were additional proteases added into the media to degrade the microgels. Thus, nanoparticle release should primarily happen inside the macrophages following microgel degradation by intracellular enzymes. Therefore, as expected, macrophages positive for nanoparticles only (in flow cytometry plots), appear at later time points, presumably after the macrophages have phagocytosed the Nano-in-Microgels and the DyLight 650 fluorescence has either degraded inside the cells or is not stable in the endosomal pH – thus shifting the flow population to the Blue Fluosphere only (i.e. nanoparticle only) flow quadrant (see Supplemental Figure 3).

**Figure 5:**
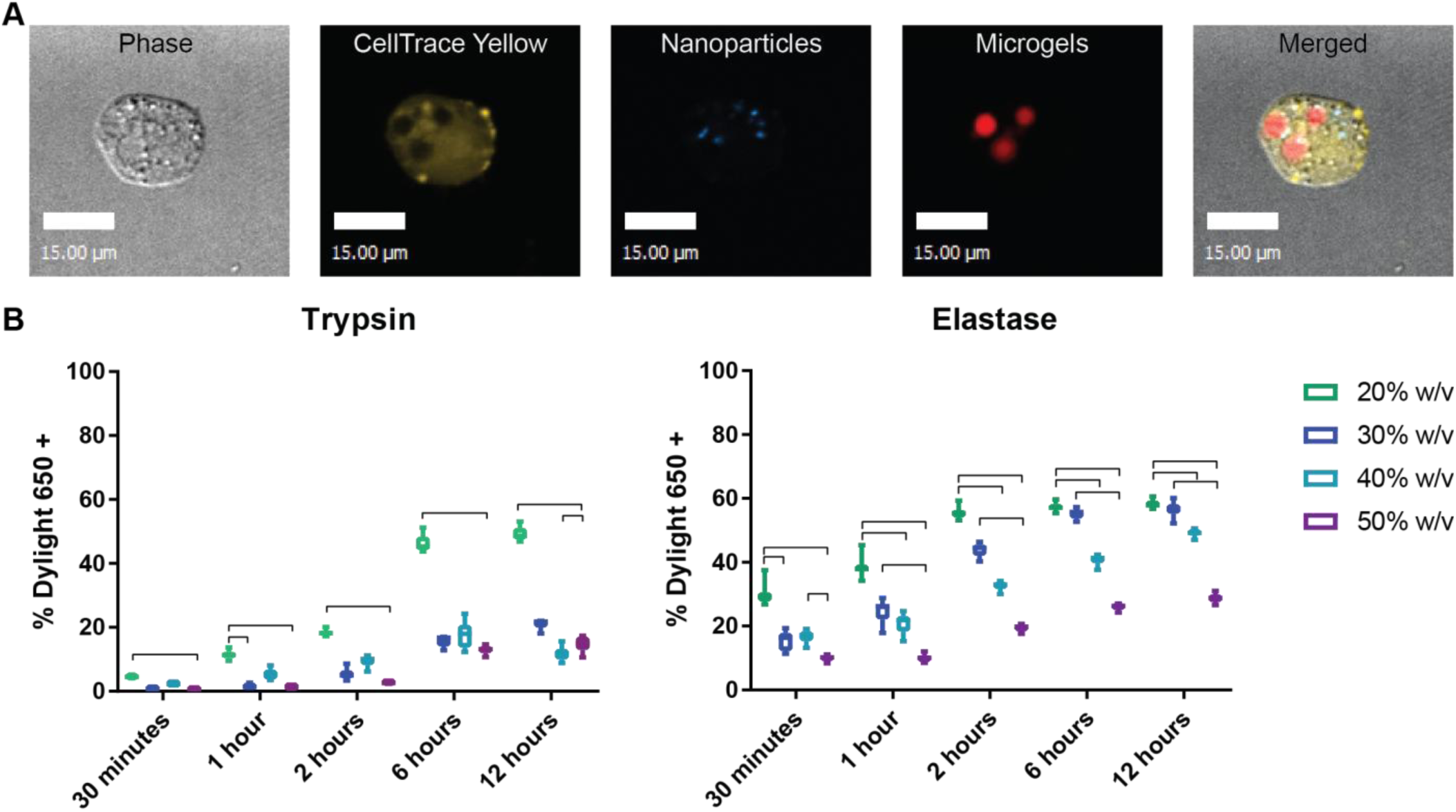
Trypsin- and elastase-responsive microgel-uptake by RAW 264.7 macrophages. Microgels were delivered at a 1:1 ratio to RAW 264.7 macrophages and run on flow cytometry to measure microgel positive macrophages at 30 minutes, 1, 2, 6, and 12 hours. (A) Representative images of a CellTrace Yellow labeled macrophage with 100 nm Blue FluoSpheres, and DyLight 650 labeled microgels; scale bar = 15 µm, images linearly contrasted for clarity. (B) Trypsin- and elastase- responsive microgel uptake over time measured by the percentage of DyLight 650 positive RAW 264.7 macrophages; trypsin- (n=6) and elastase-responsive (n=12), Kruskal-Wallis with Dunn’s multiple comparison test, bars represent statistical significance, α < 0.01.

### Microgels avoid rapid clearance following intratracheal delivery and are detectable up to 24h in naïve mice

The final design criteria was to avoid rapid phagocytic clearance by alveolar macrophages, a major barrier in pulmonary drug delivery. We chose the formulation with the fastest phagocytosis *in vitro* (20% w/v, Figure 5) to evaluate what the shortest residence time of these formulations would be *in vivo*. To understand the baseline clearance by phagocytosis or the mucociliary elevator, we chose to evaluate this in naïve animals in the absence of any disease-specific proteases which would otherwise convolute the results. We also tested clearance with the trypsin-responsive microgel formulations as trypsin is a protease prmarily found in the GI system – and not in the lungs.

Specifically, Balb/c mice were treated with saline or 20% w/v trypsin-responsive DyLight 650-mal Microgels by a Penn-Century MicroSprayer aerosolizer. Lungs were harvested within 30 minutes after microgel instillation, and processed for crysectioning. Histologically, we observed that microgels were present in all 5 lobes of the lungs. Few sections were also stained for Siglec F, an aleveloar macrophage and eosinophil marker. Our results show that some microgels are associated with SiglecF+ cells immediately after delivery (Supplemental Figure 4).

For longitudinal evaluation of microgel retention two Nano-in-Micro formulations were tested, a 20% w/v trypsin-responsive and a 20% w/v non-protease responsive PEG-PEG Nano-in-Microgel formulation; both formulations had the microgels labeled with DyLight 650 and encapsulated 100 nm Blue FluoSpheres (Blue FS). These Nano-in-Micro formulations were instilled naïve Balb/c lungs which were excised at 0.5, 6, and 24 hours, and the radiant efficiency of the whole lung was measured using IVIS. Fluorescence intensity decreased with time but was still visible at 6 and 24 hours after injection suggesting a slower clearance rate (Figure 6A-B and Supplemental Figure 5A). We expect that the levels of proteases present in these naïve lungs were relatively low, and thus the measured decrease in fluorescence should not be due to enzymatic degradation of the peptide backbone on the microgels, which was confirmed by the similar rates of clearance for both the protease- and non-protease degradable (PEG-PEG) Nano-in-Microgel formulations.

**Figure 6:**
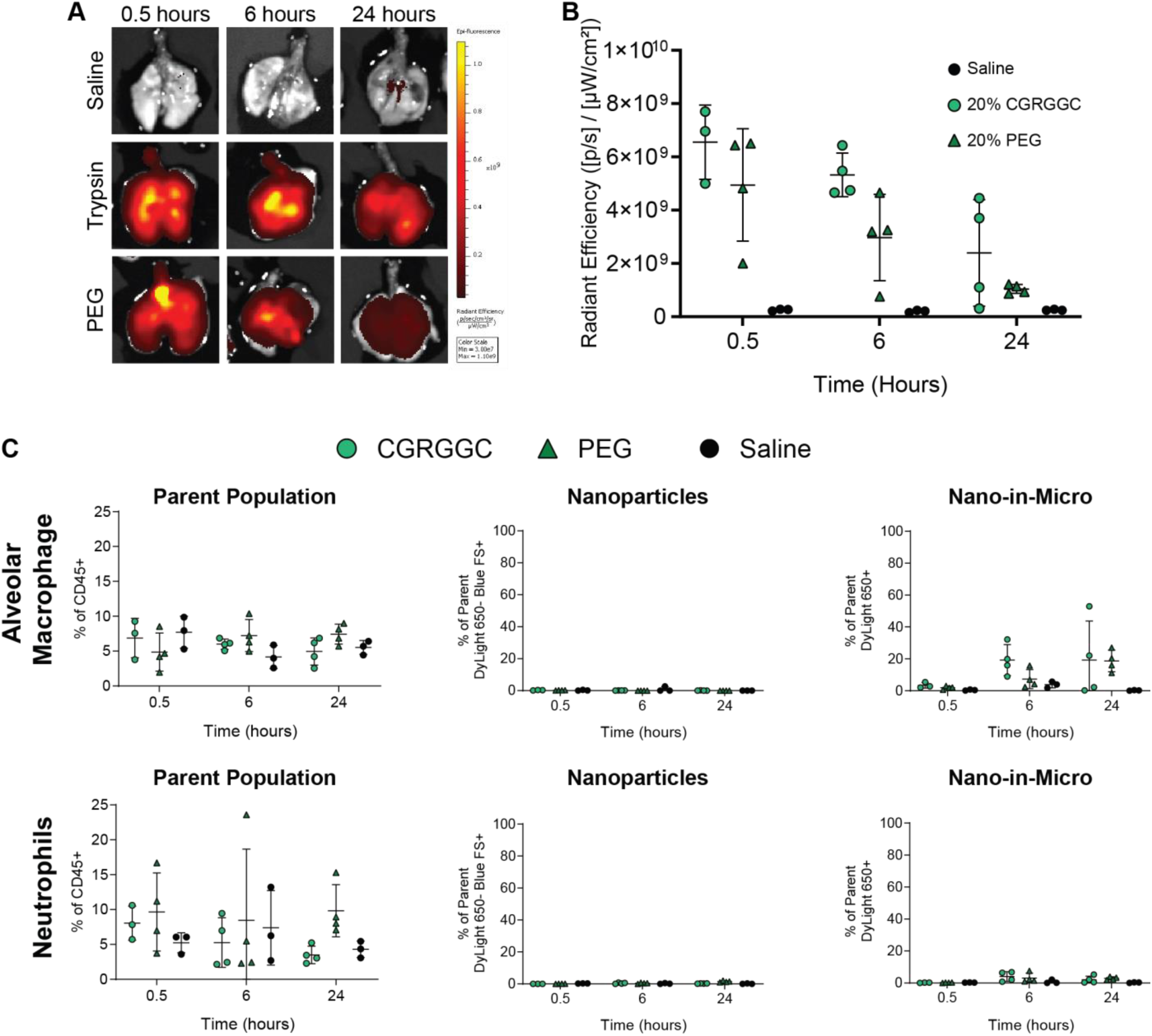
Trypsin- and non-protease(PEG)-degradable microgel residence time in naïve mice. Saline, DyLight 650 trypsin-responsive microgels, or DyLight 650 PEG (non-protease responsive) microgels were instilled in naïve mice airways using a MicroSprayer then lungs were excised and radiant efficiency measured by IVIS at 0.5, 6, and 24 hours post delivery. (A) Representative IVIS images for lung background control (saline), and DyLight 650 emission from trypsin-responsive and PEG-crosslinked microgels at 0.5, 6, and 24 hours after instillation. (B) IVIS measured radiant efficiency decreased for both formulations over time with similar rates (n = 3 - 4). (C) Flow cytometry for Alveolar Macrophages and Neutrophil populations showed no significant changes in cell populations across treatments and no uptake of free nanoparticle (DyLight 650-Blue FS+) within the macrophage or neutrophil populations across all timepoints. Limited Nano-in-Micro (DyLight 650+) uptake in the macrophages is seen after 6 hours within some individual mice but no group significance. Kruskal-Wallis with Dunn’s multiple comparison test within each time point is not statistically significant, α < 0.01.

A secondary goal of these experiments was to ensure that nanoparticles are not being released in normal, disease-free lungs – a key criterion to minimize side effects. To evaluate this we looked for the presence of the 100 nm Blue FluoSphere (ex/em 350/440; DAPI equivalent) in various cell types in the lungs of naïve mice following delivery using flow cytomtery. In the absence of significant proteolytic degradation (in naïve animals we do not expect disease-related proteolytic enzymes to be present in the lung), few Blue FluoSpheres should be released from the microgels (labeled with DyLight 650). This was confirmed by flow cytometry showing that alveolar macrophages (Live CD45+Ly6G-SiglecF+CD11c+CD11b-) and neutrophils (Live CD45+Ly6G+) were not positive for free nanoparticles (i.e. nanoparticles released out of microgels; 100 nm Blue FluoSphere+ and DyLight 650-), Figure 6C. Additionally epithelial cells (Live CD45-EpCAM+) were also negative for free nanoparticles (Supplemental Figure 5B). Importantly this also showed little to no uptake by neutrophils and minimal uptake of the Nano-in-Microgel by alveolar macrophages (lack of macrophages positive for DyLight 650) suggesting avoidance of rapid macrophage phagocytosis within the early timepoint.

Without degradation the entrapped nanoparticles or partially degraded microgels are not expected to be found in other organs, except potentially in lung draining lymph nodes (carried by lung phagocytic cells). However, IVIS images showed no detectable radiant efficiency in the lymph nodes (Supplemental Figure 5C), indicating minimal or no transport of the microgels outside of the lung. This suggests that without relevant proteases to degrade the microgels, and with minimal macrophage-phagocytosis as discussed above, they are most likely cleared by normal mucociliary clearance of the lung.

The radiant efficiency observed in 20 and 30% w/v formulations were similar (Supplemental Figure 6A-B), clearance also remained similar between the formulations, and additional flow cytometry showed no alveolar macrophages or neutrophils positive for either the Nano-in-Microgel formulations or the nanoparticles alone (Supplemental Figure 6C) suggesting limited rapid clearance of the Nano-in-Micro system for the final design criterion. Fluorescence of the Nano-in-Microgel formulation was maintained at 6 hours, showing that microgels avoid rapid clearance by phagocytosis and when designed to be used in diseased states should degrade within this time frame (< 6 h) leading to effective release of the nanoparticles.

### Limited phagocytosis *in vivo* within the first hour of administration

To show that the third design criterion was not dependent to a specific delivery method, or a specific proteolytically-responsive peptide, or a mouse strain, a 1:1 ratio of trypsin-responsive microgels and elastase-responsive microgels (labeled with DyLight 650-4xPEG mal and DyLight 488-mal, respectively) loaded with 100 nm Blue FluoSpheres were co-delivered to the airways of Balb/c or C57BL6 naïve mice. Microgels were administered with the aerosolizer (PC) or by the commonly used oropharyngeal aspiration method between Balb/c littermates. Additionally, microgels were delivered by oropharyngeal aspiration to C57BL/6 mice from two age group (5-6 and 16-17 week old) since this strain and delivery method is often used in LPS models of neutrophilic infiltration. One hour after delivery (Figure 7) the mice were sacrificed, the lungs collected, and digested into single cell suspensions. Stained samples were then run on a flow cytometer to determine the phagocytic cells positive for the different microgel formulations. Cell populations were defined as follows: 1) polymorphonuclear neutrophils (PMN): CD45+Ly6G+, 2) alveolar macrophages (AM): CD45+Ly6G-SiglecF+CD11c+CD11b-, 3) eosinophils (Eo): CD45+Ly6G-SiglecF+CD11c-CD11b+, and 4) non-alveolar F4/80+ macrophages (Mφ): CD45+Ly6G-SiglecF-F4/80+. The MicroSprayer appears to deliver microgels in the vicinity of eosinophils and alveolar macrophages, Supplemental Figure 4; however, the extent and dose with which these particles reach the lower airways could be low, and with oropharyngeal aspirations may be even lower since the velocity added by the MicroSprayer is absent, this could result in lower percentages of particle uptake. Additionally, the F4/80+ macrophage populations can include both non-alveolar macrophages as well as interstitial macrophages [38] that may or may not come in contact with the microgel formulations, also skewing the percentages low.

**Figure 7:**
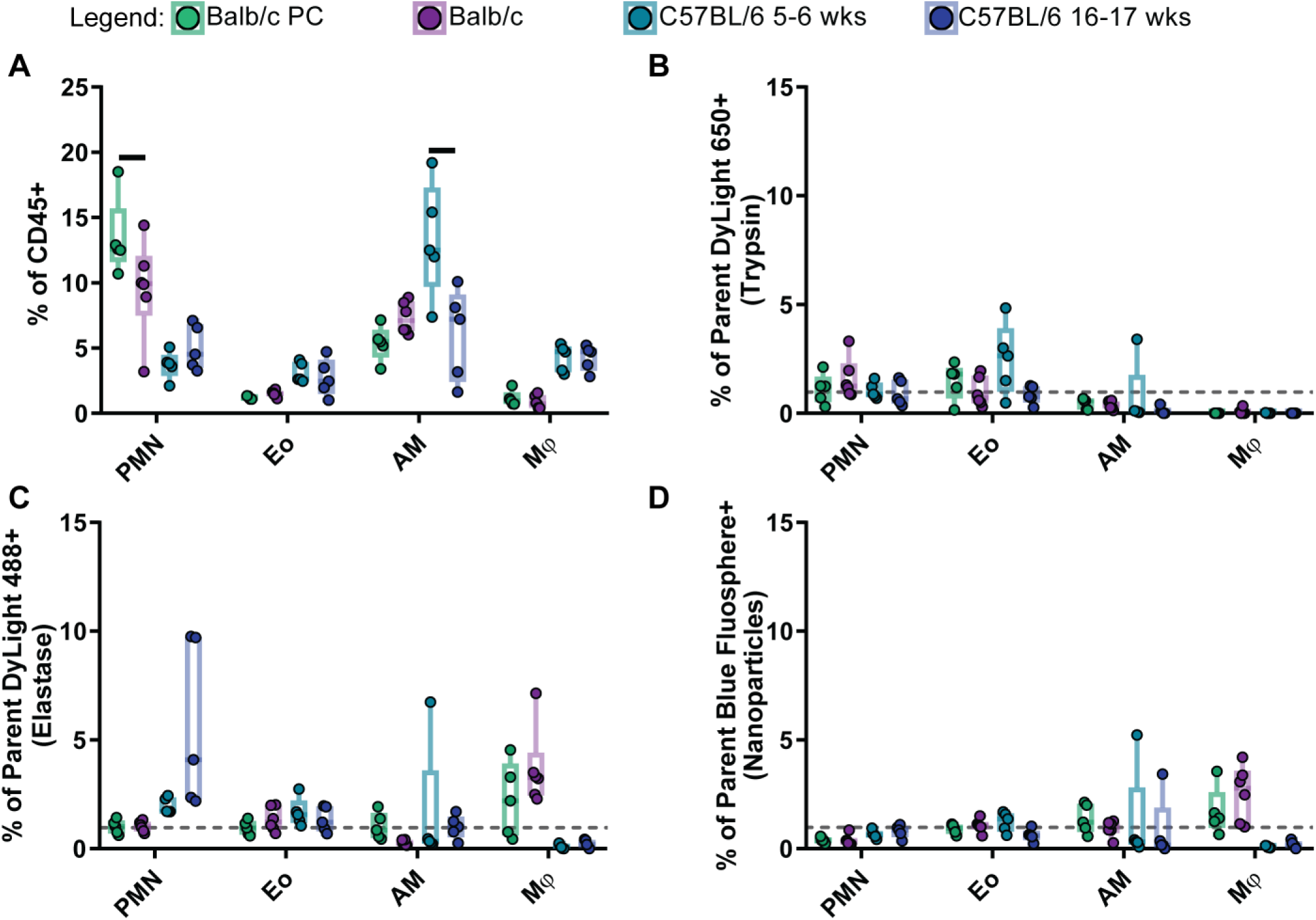
20% w/v trypsin- and elastase-responsive microgel uptake in naïve Balb/c or C57BL6 phagocytic populations one hour post-delivery. 20% w/v DyLight 650 trypsin-responsive microgels encapsulating 100 nm Blue FluoSpheres and 20% w/v DyLight 488 elastase-responsive microgels encapsulating 100 nm Blue FluoSpheres were instilled at a 1:1 ratio in mice airways by either Penn-Century MicroSprayer (PC) or oropharyngeal aspirations in naïve Balb/c littermates (PC-green, purple) or by oropharyngeal aspirations to 5-6 week-old (light blue) or 16-17 week-old (dark blue) C57BL/6 naïve mice. Single cell suspensions were stained and run on flow cytometry (A) Percentage of neutrophils (PMN; CD45+Ly6G+), eosinophils (EO; CD45+Ly6G-SiglecF+CD11c-CD11b+), alveolar macrophage (AM; CD45+Ly6G-SiglecF+CD11c+CD11b-), and F4/80+ macrophage (Mφ; CD45+Ly6G-SiglecF-F4/80+) of the CD45+ population; two-way Anova (α = 0.01) with Sidak post hoc. Percentage of neutrophils, eosinophils, alveolar macrophages, and F4/80 macrophages positive for (B) trypsin-responsive microgels (DyLight 650+) or (C) elastase-responsive microgels (DyLight 488+) or (D) Blue FluoSpheres (ex/em 350/440; DAPI channel); Mann-Whitney (α = 0.01). Grey dashed line marks the 1% threshold used to set the gates on FMOs.

The neutrophil populations for the two Balb/c delivery methods were significantly different and the alveolar macrophage population between the two C57BL/6 groups were also significantly different using a two-way Anova (α = 0.01) with Sidak post hoc test, Figure 7A. Statistical analysis comparisons were kept within species for each cell type. While the microgel+ populations within each of the cell populations have median values below 5% for each type, (Figure 7B-C), the elastase-responsive group has a few between 5-10%; the grey dashed line marks the 1% threshold used to set the gates on FMOs. This data suggests limited rapid phagocytosis within this short timepoint, achieving the final design criterion. Due to the low number of cells within each of the sub-populations the nanoparticles (Figure 7D) were gated directly off the AM, PMN, Eo, or Mφ cell population rather than from the microgel (DyLight 650 or DyLight 488) positive populations with only the Balb/c F4/80+ macrophage populations showing cells positive for nanoparticles.

While particle size is an important factor for phagocytosis and endocytosis, recent work looking at the relationship between particle elasticity and cellular uptake has shown that softer particles are often more difficult for cells to consume. This is because particle deformation leads to a greater extent of spreading between the elastic particle and the cell; specifically, phagocytosis by immune cells for particles in the softer (kPa) range having slower uptake than stiffer (MPa) particles[37, 39]. We observed that with the 20% w/v trypsin- and elastase-responsive microgels, there was minimal rapid clearance by phagocytosis in the first hour by neutrophils, eosinophils, alveolar macrophages, and F4/80+ macrophages for both Balb/c and C57BL/6 mice regardless of delivery method (aerosolizer vs. oropharyngeal aspiration); we believe the larger size of the microgels in the swelled state and their low mechanical stiffness leads to lower phagocytic clearance.

Furthermore, that *in vitro* elastase-responsive microgels degraded in CF patient sputum within 30 minutes, which suggested that the elastase-responsive microgels should degrade and release nanoparticles before most of the microgels are cleared from diseased lungs. This microgel phagocytosis and clearance from Figures 6-7 was observed in naïve animals, however, under severe disease burdens (such as asthma, COPD, and CF) there is reduced phagocytic ability of these innate cells[40–43] which may create a longer window for the microgels to degrade—releasing their cargo.

## Conclusions

The Nano-in-Microgel system was designed for (1) appropriate aerodynamic size to ensure deep lung delivery (2) protease-triggered release of encapsulated nanoparticles in diseased lung microenvironments and (3) avoiding rapid clearance by alveolar macrophages. The purpose of these studies was to understand two issues: 1) how the starting polymer concentration of our Nano-in-Microgel multistage particles influenced the size and degradation *in vitro* and 2) the cellular-biodistribution, phagocytosis, and clearance of the Nano-in-Microgel system in naïve *in vivo* settings following pulmonary delivery. The trypsin peptide crosslinker was used to understand how the Nano-in-Microgel system without specific enzymatic degradation cleared *in vivo*, while the neutrophil-elastase responsive microgels demonstrated that we could incorporate a pulmonary disease-relevant crosslinker into the system and study nanoparticle release. We have further improved the MADE method by changing the chemistry from an acrylate-thiol to maleimide-thiol for faster reaction kinetics and lowering the necessary % w/v for the microgel formulations while maintaining the size ranges needed for deposition and avoiding rapid clearance by phagocytosis within the first 24 hours. While the work used representative fluorescent nanoparticles, these nanoparticles can be easily exchanged for target-specific nanoparticles as they are physically entrapped in the microgel network during fabrication.

Future studies with the elastase-responsive peptide in disease models with therapeutic nanoparticles should be completed to determine this optimal clearance vs. degradation kinetics and to understand the efficacy of the Nano-in-Microgel system for drug delivery under various disease conditions.

The results of these studies show that the MADE method for fabricating multi-stage particles produces enzyme-responsive microgels without exposure to degradative conditions for biologics that allow for quick release of nanoparticles for targeting pulmonary disease. Future work in optimizing a dry powder formulation for existing pulmonary delivery devices (e.g., DPIs, nebulizers) is needed to translate the system into clinical usage.

## Acknowledgements

This work was supported by funding from the Marcus Center for Therapeutic Cell Characterization and Manufacturing (MC3M) at Georgia Tech, Georgia Tech Foundation (KR), Georgia Tech Research Alliance (KR), Center for Cystic Fibrosis Airways Disease Research and Children’s Healthcare of Atlanta, NIH/NIGMS-sponsored Cell and Tissue Engineering (CTEng) Biotechnology Training Program T32GM008433 (JCM) and NSF Graduate Research Fellowship DGE-1148903 (JCM). The authors would like to thank the Prof. Hang Lu and her lab for the use of their cell traps, Prof. Andrés García’s lab for the use of their Rheometer, Dr. Randall Toy for help during animal experiments, and Prof. Rabin Tirouvanziam for help with Emory’s Cystic Fibrosis Biospecimen Registry. We also thank the Parker H. Petit Institute for Bioengineering and Bioscience at the Georgia Institute of Technology for the use of their core facilities.

## Supplemental Data

Supplementary data related to this article can be found online

