## Supplemental for "*In-vitro* and *in-vivo* Characterization of a Multi-Stage Enzyme-Responsive Nanoparticle-in-Microgel Pulmonary Drug Delivery System"

Graphical Abstract

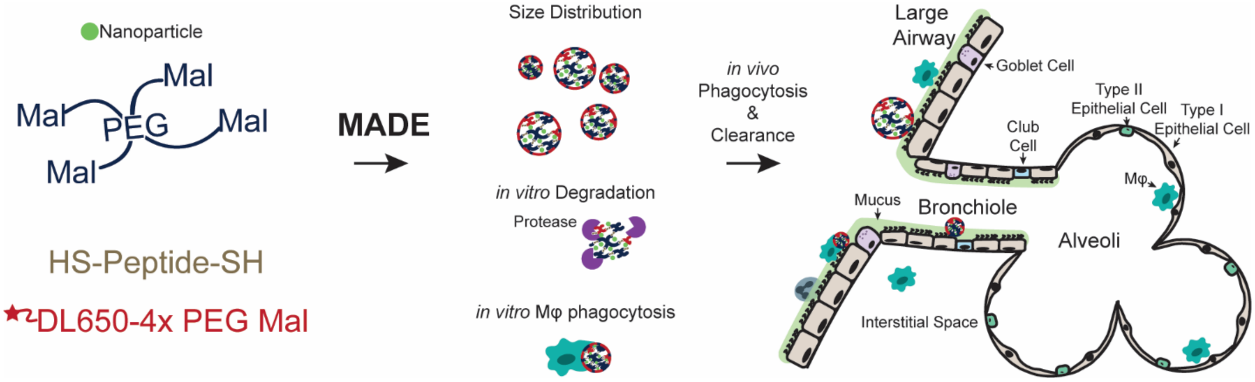


| Supplemental Table 1: Microgel Formulations | | | | | | |
|---|---|---|---|---|---|---|
| | | | Trypsin Responsive | | Elastase Responsive | |
| | | | Peptide | PEG-4-Mal | Peptide | PEG-4-Mal |
| **w/v%** | **Vf (µL)** | **W (mg)** | (mg) | (mg) | (mg) | (mg) |
| 60 | 100 | 60 | 6.6 | 53.4 | 12.0 | 48.0 |
| 50 | 100 | 50 | 5.5 | 44.5 | 10.0 | 40.0 |
| 40 | 100 | 40 | 4.4 | 35.6 | 8.0 | 32.0 |
| 30 | 100 | 30 | 3.3 | 26.7 | 6.0 | 24.0 |
| 20 | 100 | 20 | 2.2 | 17.8 | 4.0 | 16.0 |

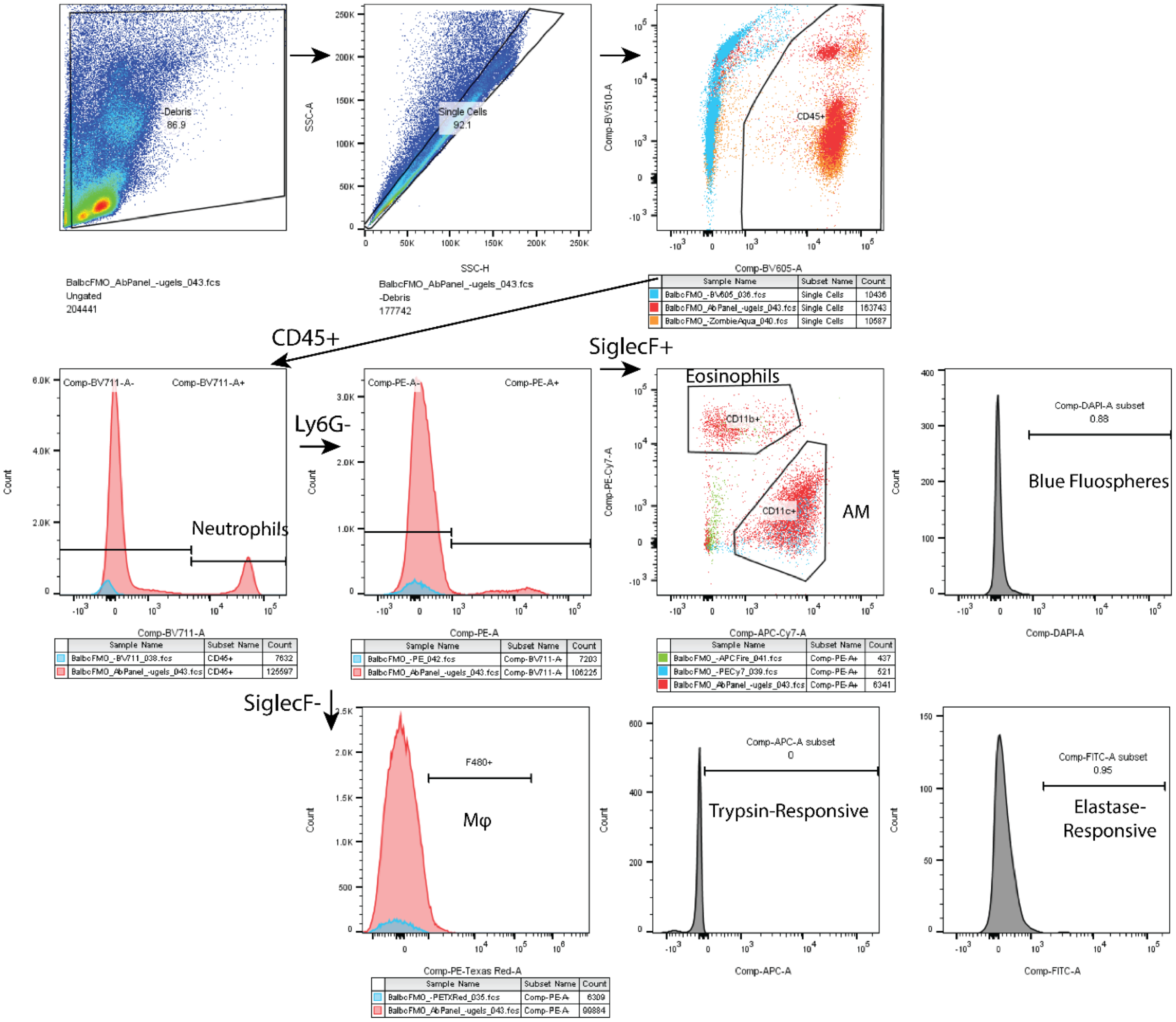


**Figure S1:** Gating scheme and FMOs for flow panel. Dead cells were excluded from analysis in S.Figure 4 and S.Figure5; however, due to the few number of cells in the final eosinophil and alveolar macrophage gates in Figure 7, the dead cells were included in the analysis of that experiment. Neutrophils were gated as CD45+ Ly6G+, Eosinophils as CD45+ Ly6G+ SiglecF+ CD11b+ CD11c-, Alveolar Macrophages as CD45+ Ly6G- SiglecF+ CD11b- CD11c+, and macrophages gated as CD45+ Ly6G+ SiglecF- F4/80+ then each population was gated for trypsin (APC)- or elastase (FITC)-responsive microgels and Blue FluoSpheres (DAPI equivalent). Blue FluoSpheres were directly gated on the cell population as the numbers were too low to gate on the negative of the trypsin- or elastase-responsive microgels.

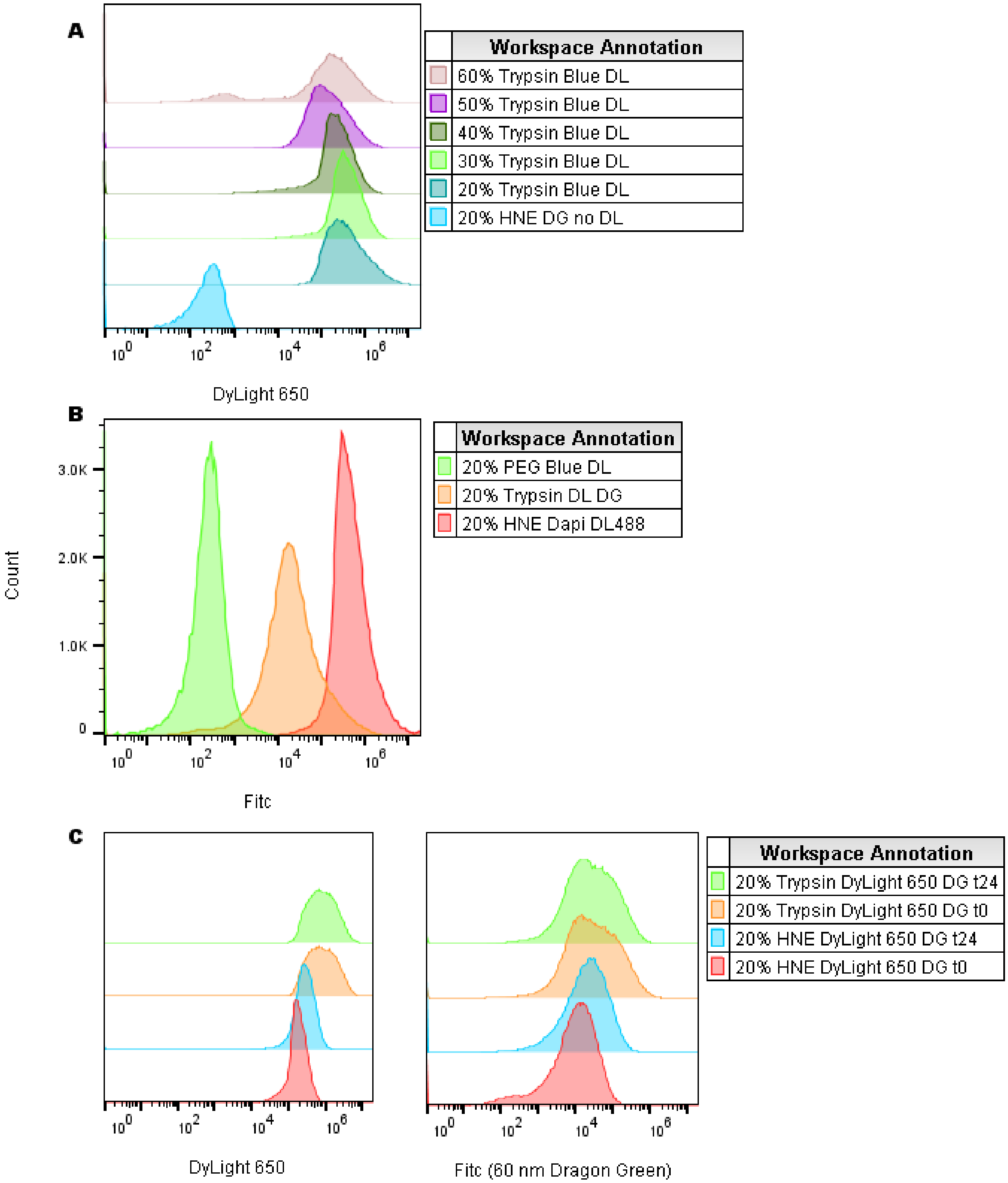


**Figure S2:** (A) Accuri signal of fluorescently DyLight 650 labeled microgels for the following formulations; no DyLight 650 (cyan), Trypsin-responsive 20% (blue), 30% (light green), 40% (dark green), 50% (purple), 60% (light purple) (B) Microgels without Dragon Green (DG) nanoparticles (light green), with 60 nm Dragon Green nanoparticles (orange), and microgels labeled with DyLight 488 (red). (C) Microgel fluorescence on Day 0 for 20% w/v Trypsin (Light Green) and 20% Elastase (red) and fluorescence 24h later for the same batch (orange, blue respectively).

| Supplemental Table 2: 100 nm Blue Fluosphere Encapsulation | | |
|---|---|---|
| % w/v | % Enapsulation | # NP Encapsulated |
| | **Trypsin (CGRGGC)** | |
| 20 | 34% ± 1% | 1.2E11 ± 1.9E9 |
| 30 | 30% ± 1% | 1.1E11 ± 3.7E9 |
| 40 | 42% ± 5% | 1.5E11 ± 1.7E10 |
| 50 | 31% ± 2% | 1.1E11 ± 6.1E9 |
| | **Elastase (CGAAPVRGGGGC)** | |
| 20 | 33% ± 1% | 1.2E11 ± 4E9 |
| 30 | 33% ± 3% | 1.2E11 ± 9.2E9 |
| 40 | 31% ± 2% | 1.1E11 ± 6.8E9 |
| 50 | 32% ± 2% | 1.14E11 ± 8E9 |

| Supplemental Table 3: Estimated Pore Size | |
|---|---|
| % w/v | nm |
| | **Trypsin (CGRGGC)** |
| 20 | 20 |
| 30 | 14 |
| 40 | 14 |
| 50 | 16 |
| | **Elastase (CGAAPVRGGGGC)** |
| 20 | 10 |
| 30 | 13 |
| 40 | 16 |
| 50 | 19 |

**Method:** Pore size was calculated using the following equations:

(1) $(\bar{r}_0^2)^{1/2} = l\left(\frac{\bar{M}_c}{M_r}\right)^{1/2} C_n^{1/2}$

(2) $v_{2,s} = \frac{V_p}{V_s} = \frac{1}{Q}$

(3) $\xi = v_{2,s}^{-1/3}(\bar{r}_0^2)^{1/2}$

For equation 1, the bond length, l (1.46 Å), molecular weight of the repeating unit, $M_r$ (44 g/mol), and characteristic ratio $C_n$ (4), were used from PEG, which may also underestimate the mesh size. The average molecular weight between cross-links, $M_c$, was calculated from the combined PEG and peptide molecular weights. The swelling ratio was calculated from the mean size in the swollen (Vs – swollen volume) and relaxed (Vp – dry volume) states measured by microscopy; this was within range of previously reported swelling ratio of the 60% w/v acrylate Trypsin microgels. This gives an estimated mesh size ranging from 13-20 nm.

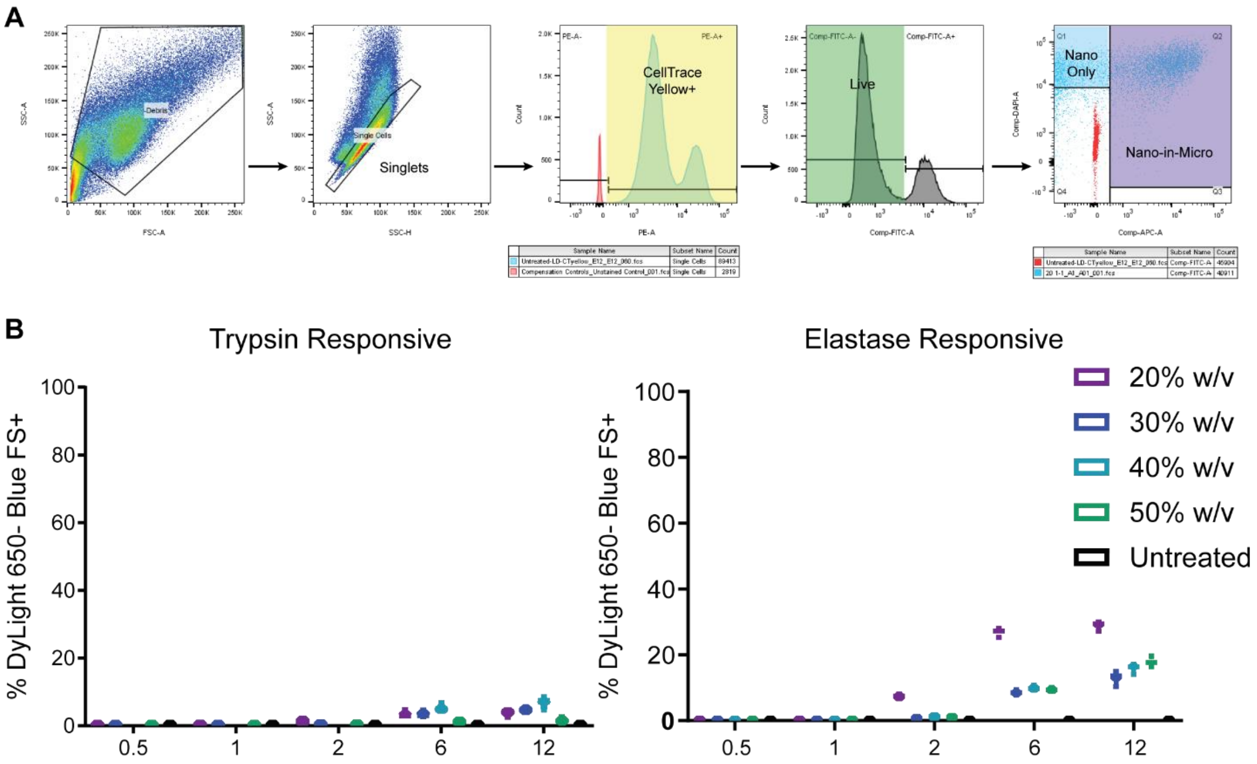


**Figure S3:** RAW 264.7 Macrophages were stained with CellTrace Yellow, given Nano-in-Microgels (100 nm Blue FluoSpheres, DyLight 650 microgels) at a 1:1 ratio, stained with Zombie Green (live/dead) and fixed at 0.5, 1, 2, 6, 12 hours (A) Flow cytometry gating schematic. Singlets were gated for the CellTrace Yellow (PE) positive cells, followed by gating on live cells (FITC-negative) then finally gated for cells containing nanoparticles only (DAPI-equivalent Blue FluoSpheres; these are nanoparticles after release from microgels extracellularly and then phagocytosed, or after microgels have been fully degraded inside the cells following Nano-in-Microgel uptake) or cells containing the intact Nano-in-Microgels (double positive DAPI and APC). (B) Flow results with a slight increase in macrophages positive at the longer time points. This increase at 6 – 12h is possibly a result of the macrophages that consumed the Nano-in-Micro formulation early and the DyLight 650 is either degraded or not stable in the lower endosomal pH. n=6 for trypsin-responsive, n=12 for elastase responsive, nanoparticle data for 40% w/v trypsin-responsive was not collected at 0.5, 1, 2h.

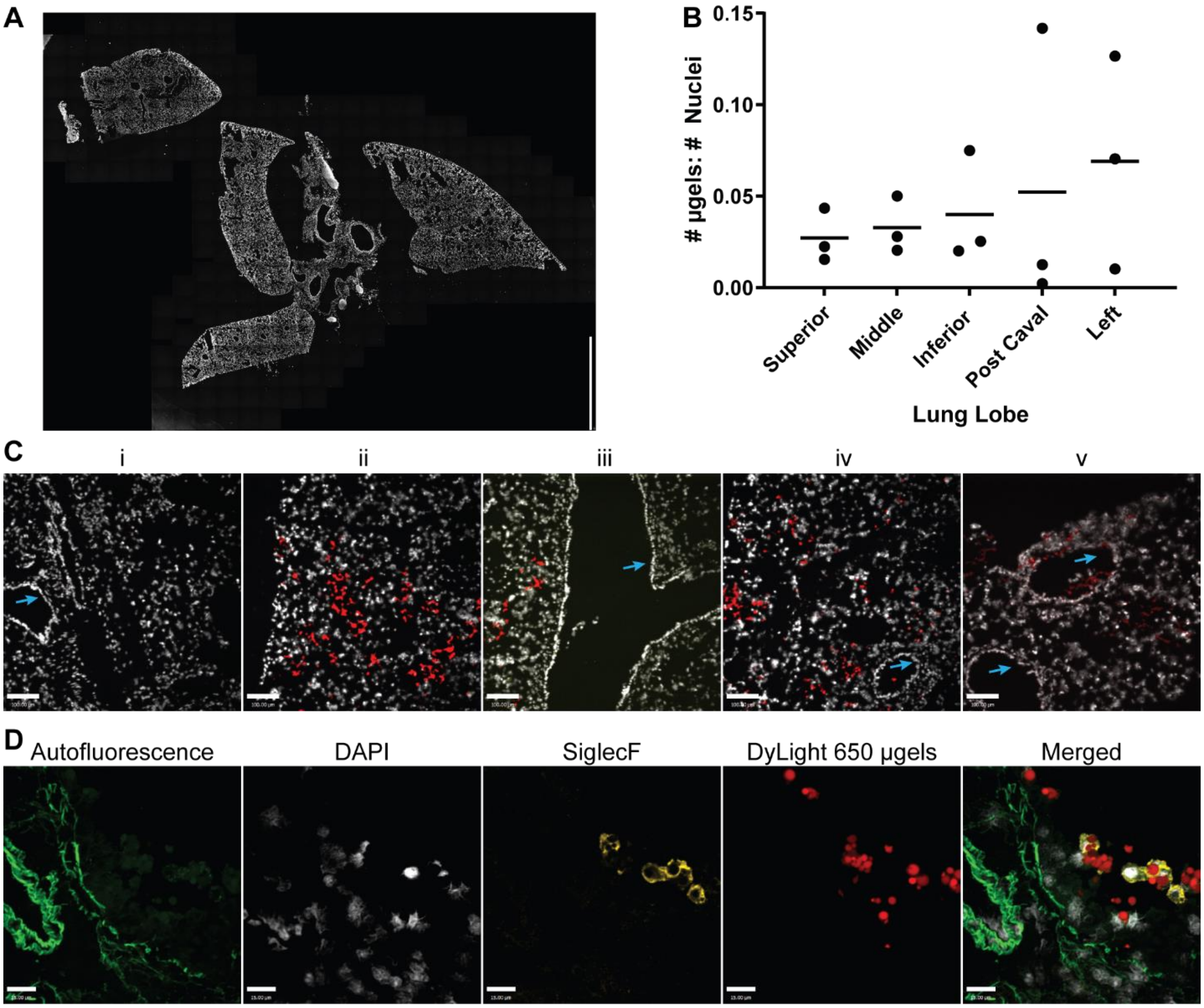


**Figure S4:** DyLight 650-mal Microgels (without fluorescent nanoparticles) were instilled in mice lungs. Lungs were harvested within 30 minutes, frozen for cryosectioning and stained with SiglecF and DAPI. (A) Representative stitched image of DAPI (white) stained whole lung section (right side scale bar = 3 mm). (B) Ratio of microgels to nuclei within each lobe, each point is an individual mouse and shows microgel deposition within each lobe. (C) Representative images of microgel (red) deposition in different areas within the sections, blue arrows point to larger vessels which have denser populations of nuclei (i, iii, iv, v). There is some variability in the distribution across the sections with some having limited microgels present (i) while others have more (ii). Microgels appear to deposit in the deep lung, however, some microgels are present in the larger airways (iv and v, blue arrows), scale bar = 100 µm. Due to the objective used and the reduced alfalfa free diet, tissue autofluorescence was not detectable at this magnification. (D) 60x imaging showing tissue autofluorescence, nuclei, SiglecF stained cells (eosinophils or alveolar macrophages), and DyLight650+ microgels within a lung section (scale bar 16 µm). Images linearly contrasted for clarity.

ImmunoFluorescence Staining: After IVIS, lungs were fixed in 4% paraformaldehyde at 4 C overnight, allowed to settle in 30% w/v sucrose then frozen in OCT in a dry ice isopentane mixture. 5-10 µm lung sections were sliced and stored at -80 C until they were processed. Slides were washed in 1x TBS for 5 minutes, placed in cold acetone for 10 minutes, then washed again in 1x TBS. For whole lung imaging slides, Prolong Gold DAPI was added, allowed to cure overnight, sealed then imaged. For Siglec-F staining, after the second TBS wash, sections were blocked for 2 hours at room temperature in 5% goat serum 0.5% Triton X in 1x TBS. Purified SiglecF at 10 µg/mL in blocking buffer was left on the sections at 4 C overnight. Slides were washed in TBS and stained with a Goat Anti Rat Alexa Fluor 555 secondary antibody 5 µg/mL for 30 minutes at 37 C.

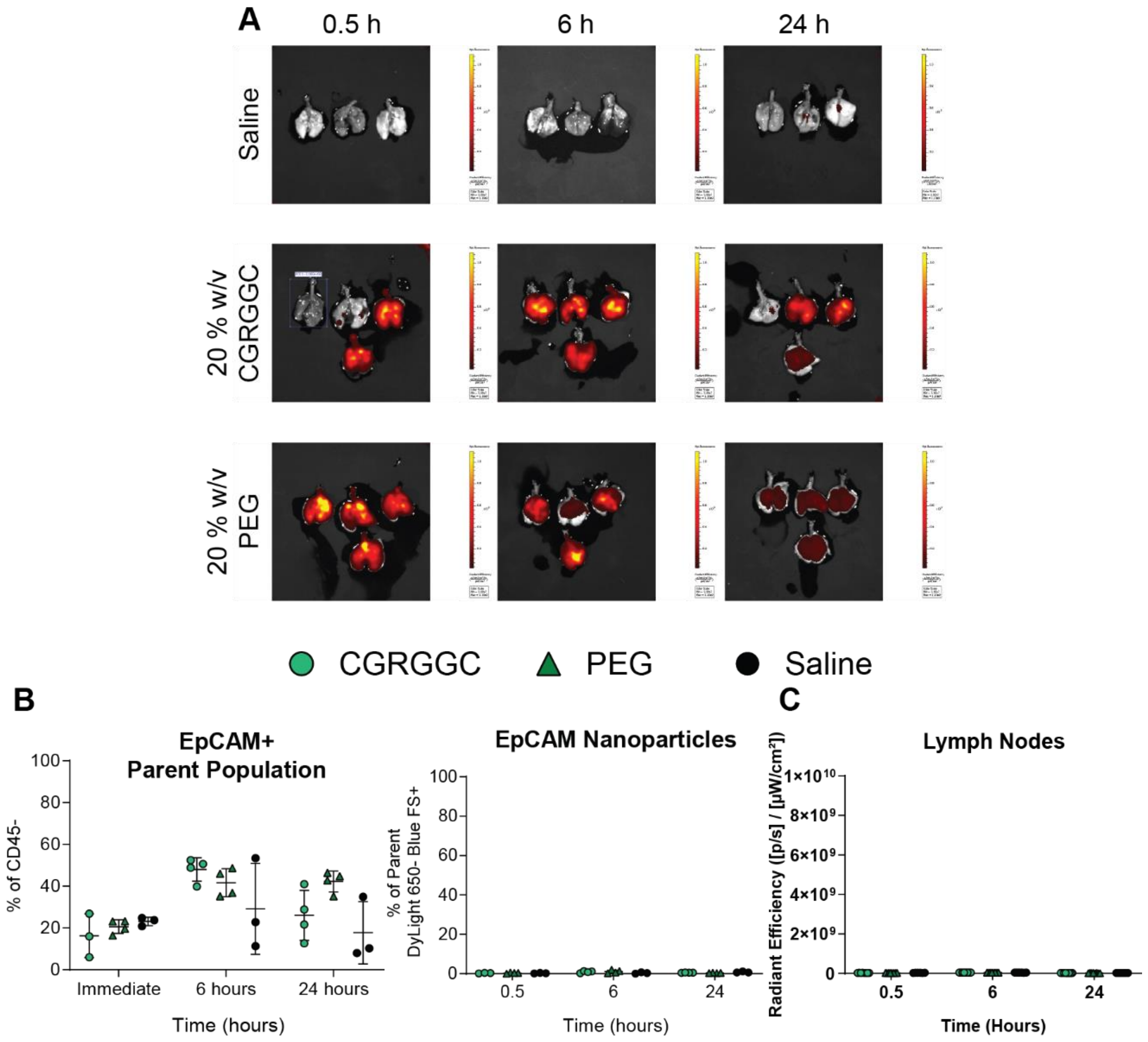


**Figure S5:** Trypsin- and non-protease(PEG)-degradable microgel residence time in naïve mice. (A) All IVIS images for 20% Trypsin and 20% PEG microgel lung delivery, the lung from 0.5h trypsin with an ROI box was a sample injected into the esophagus and removed from the data sets. (B) Flow cytometry of epithelial cells gated as CD45-EpCAM+ are negative for nanoparticles only (DyLight 650- Blue FS+). (C) IVIS radiant efficiency for the lung draining lymph nodes was not positive for microgels (DyLight 650).

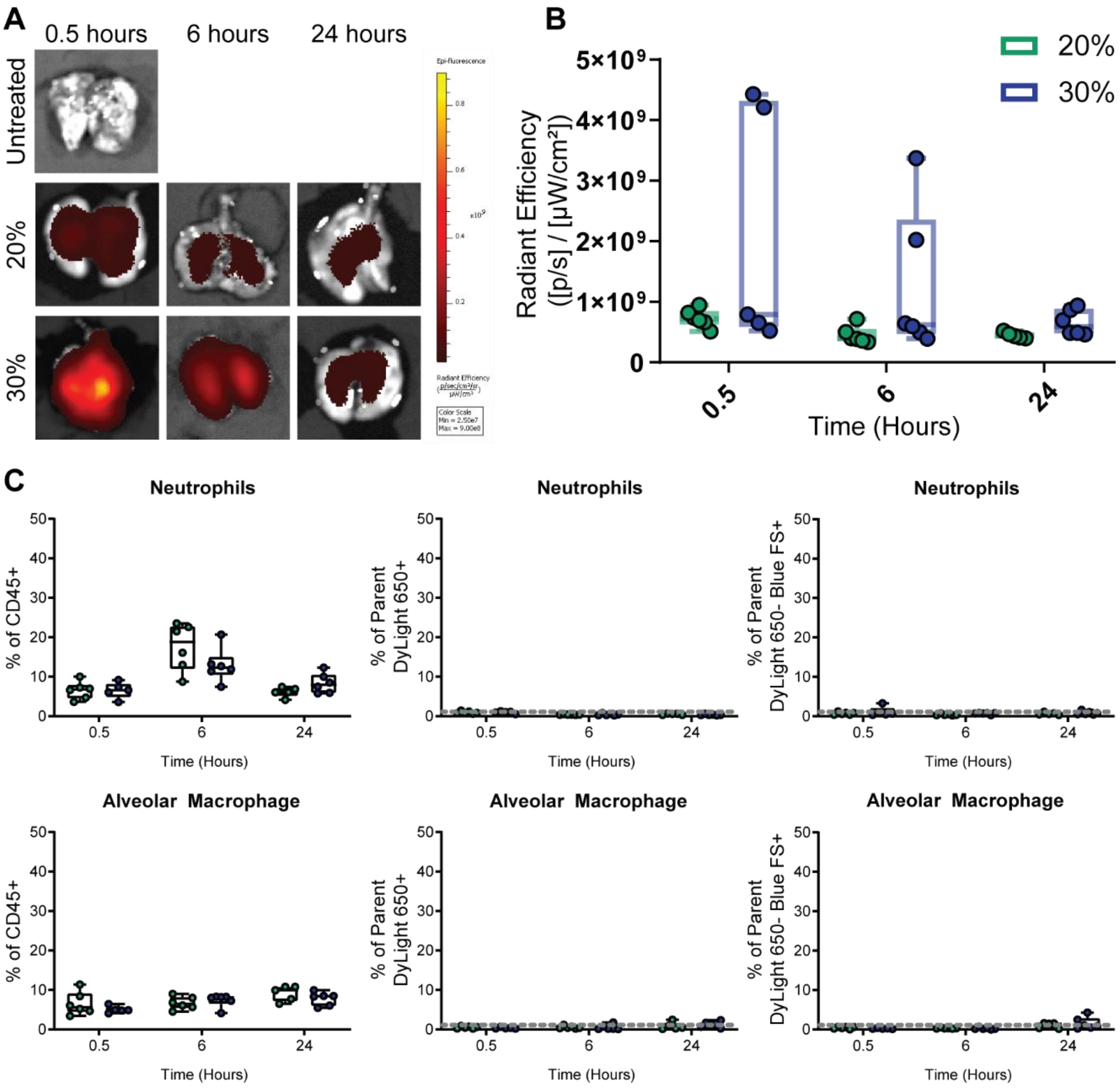


**Figure S6**: 20 and 30% w/v trypsin-responsive microgels clearance in naïve mice. DyLight 650 trypsin-responsive microgels fabricated with a starting polymer concentration of 20 or 30% w/v carrying 100 nm Blue FluoSpheres were instilled in naïve mice airways using a MicroSprayer then lungs were excised and radiant efficiency measured by IVIS at 0.5, 6, and 24 hours post-delivery (A) Representative IVIS images for radiant background efficiency (untreated), 20 and 30% w/v trypsin-responsive microgels. (B) Measured radiant efficiency of delivered 20% or 30% w/v microgel fluorescence in the excised lungs, n = 5-6 per group. (C) Flow cytometry results of the single cell lysate showing no uptake of the Nano-in-Microgels (DyLight 650+) or nanoparticles alone (DyLight 650- Blue FS+) in either neutrophils or alveolar macrophages (grey dashed line = FMO).
